# Urinary Metabolomic Profile is Minimally Impacted by Common Storage Conditions and Additives

**DOI:** 10.1101/2024.02.12.579937

**Authors:** Kelly C. Weldon, Morgan Panitchpakdi, Andrés Mauricio Caraballo-Rodríguez, Alan J. Wolfe, Pieter C. Dorrestein, Linda Brubaker, Lindsey A. Burnett

## Abstract

Metabolomics reflects the molecular communications within a biological system. Urine can be obtained non-invasively and is rich in metabolites that are potential biomarkers for human health and disease. The optimal urine storage conditions for metabolomic analysis are unknown. We measured the impact of common storage conditions and the presence of a DNA-stabilizer, AssayAssure® (Thermo Scientific), on metabolite content of voided human urine. Urinary metabolite composition was not altered across storage conditions. Although addition of AssayAssure® significantly alters metabolic profile by adduct addition, this does not preclude the identification of parent metabolites or overshadow biological differences. These data suggest biobanked urine stored in any of the tested conditions for metabolomic studies may be considered for metabolomic analysis.

## Introduction

The discovery of novel biomarkers is critical to understanding disease states and progression^1^. Metabolomics is the study of metabolic profiles in biological systems and represents the cumulative interaction of genetic, physiologic, and environmental interactions^2,3^. Unlike many biofluids, urine can be obtained non-invasively, is rich in metabolites, and reflects systemic alterations in human health and disease^4^. An important consequence of these properties is that urine is suitable for the study of health and disease and urinary metabolomics has the potential to discover novel biomarkers associated with disease states.

There are important considerations for collection and storage of urine for metabolic analysis. Metabolites can be readily altered by time and storage temperature and these alterations can contribute to discrepancies in samples collected under differential conditions^5–8^. In ideal circumstances, urine for metabolomic analysis would be snap frozen and analyzed rapidly. Unfortunately, in most clinical settings this is neither feasible nor practical.

In addition, many of the existing urine biobanks were collected and designed for other applications, such as urinary microbiome research, with the addition of preservatives and additives for those types of analysis. To date, it is unknown how the urine metabolome is impacted by commonly used additives used for other analyses such as DNA sequencing.

To determine how urine metabolite content is impacted by storage conditions and additives, we obtained voided urine and systematically investigated four commonly encountered storage conditions: refrigeration (2-4 h), simulated shipping, frozen at -80 (3 months) and compared this to quick frozen urine. All conditions were also replicated in the presence of the preservative AssayAssure®. All samples were then subjected to untargeted metabolomic analysis. The objective of our study was to compare the impact of storage and AssayAssure® addition on the metabolite content of voided urine.

## Materials and Methods

### Study Participants and Urine Sample Collection

After IRB approval, 10 adult women 18 years or older at the University of California San Diego Health’s Women’s Pelvic Medicine Center were invited to participate in this IRB-approved study (Protocol Number: 801735). Informed research consent was obtained from all participants. Women with known anatomic abnormalities of the urogenital tract, with neurologic or immunologic disease, with a history of bladder malignancy, current systemic infection, or known pregnancy were excluded. All participants contributed a midstream, voided urine specimen, which was collected using the standard “clean catch” protocol. Participants were instructed to wash their hands with soap and water, cleanse the peri-urethral area using a sterilizing wipe, discard the initial urine stream, and then collect midstream urine into a sterile Becton Dickinson (BD) Vacutainer cup.

### Urine Sample Preparation

200μl volume aliquots of midstream voided urines were immediately pipetted into microcentrifuge tubes in triplicate and subjected to the following storage conditions: 1) snap frozen in ethanol bath on dry-ice, 2) 4C refrigeration for 2-4 hours prior to extraction, 3) transport on wet ice and left in insulated box for 24 hours prior to extraction to simulate postal shipping methods prior to extraction, and 4) transport on dry ice to the lab followed by storage in -80 for three months prior to extraction. AssayAssure^®^ (Thermo Scientific, SKU 14401) was added to half of the samples prior to simulated storage.

### Metabolomics Sample Preparation

At designated timepoints sample aliquots were extracted by adding 100% MeOH with 1μM of sulfamethazine as a synthetic internal standard, to achieve a final concentration of 80% MeOH. The samples were vortexed then incubated at -20 °C for 20 minutes to aid in protein precipitation. The samples were then centrifuged for 15 minutes at 14,000 rpm and 80% of the supernatant 800μL was transferred to a prelabeled 96-Well DeepWell plate. The plates were concentrated using a CentriVap Benchtop Vacuum Concentrator until dry. Plates were covered and stored at -80 °C until data acquisition.

For data acquisition the plates were resuspended in 250μL of a 50% MeOH to water solution containing 1μM of sulfadimethoxine as an internal standard. Untargeted metabolomics data were collected using ultra-high-performance liquid chromatography system (Vanquish, Thermo Fisher Scientific, Waltham, MA, USA) with a Kinetex 1.7 um 100 A pore size C18 reversed phase UHPLC column 50 × 2.1 mm (Phenomenex, Torrance, CA, USA) coupled to an orbitrap mass spectrometer (Q-Exactive Hybrid Quadrupole-Orbitrap, Thermo Fisher Scientific, USA). The mobile phase solvent A was water with 0.1% formic acid, mobile phase solvent B is acetonitrile with 0.1% formic acid (LC-MS grade solvents, Fisher Chemical, USA). The flow rate was set to 0.500 mL/min and a 10 min per sample run time was used with a linear gradient as follows: 0 to 1 min, constant at 5% B; 1 to 7 min, linear increase to 99% B; 7 to 8 min, constant at 99% B; 8 to 8.5 min, linear decrease to 5% B; 8.5 to 10 min, constant at 5% B. Positive mode electrospray ionization was used.

### Metabolomics Data Processing and Analysis

Data in the .raw format was converted to .mzXML files using the ProteoWizard tool MSConvert. All data (.raw and .mzXML) was uploaded to MassIVE, saved under the ID MSV000091929 (ftp://massive.ucsd.edu/MSV000091929/). MS1 feature finding was conducted using the GNPS LC MZmine2 workflow, set to the preexisting batch mode: “Thermo QExactive – Kelly SEED – Mzmine-2.53)” and MZmine2 version 2.53. Three MZmine jobs were conducted for this study with these parameters: a first job including all samples, blanks and QC samples, a second job including just the samples contained in the AssayAssure® solution, and a third job of just the samples not stored in AssayAssure®. Three jobs were conducted as the presence of the AssayAssure® altered the parameters of MZmine, therefore, to understand the metabolite presence in samples without AssayAssure®, the separate jobs were necessary. All three MZmine feature tables, along with sample metadata, were then input into the global natural products social (GNPS) feature based molecular networking (FBMN) tool^9^ for molecular networking and library ID generation, using all publicly available libraries and suspect libraries^10^. The .qza files produced from GNPS and sample metadata were used as inputs for QIIME 2^11^ to perform the beta diversity analysis of the data, principal component analysis (PCoA) and biplots, and random forest analyses (https://docs.qiime2.org/2023.9/). Finally, the .graphml files were downloaded from the GNPS FBMN jobs and input into the free Cytoscape^12^ software in order to visual the molecular networks.

### Statistical Analysis

For global metabolome comparisons, PERMANOVA calculations were performed on Bray Curtis distance matrices using QIIME 2 with p<0.05 as significant. For comparison of relative abundance of individual metabolites, Kruskal-Wallis nonparametric analysis followed by Dunn’s multiple comparison tests were performed with p<0.05 as significant.

## Data Citation and Availability

UCSD MassIVE ID: MSV000091929

ftp download: ftp://massive.ucsd.edu/v07/MSV000091929/

Full dataset: https://gnps.ucsd.edu/ProteoSAFe/status.jsp?task=d434e978d972449b89eeeefd7c8d1816

AssayAssure® Dataset: https://gnps.ucsd.edu/ProteoSAFe/status.jsp?task=4366f3b00e7c44219affec6f1dd0f91e

Non AssayAssure® Dataset: https://gnps.ucsd.edu/ProteoSAFe/status.jsp?task=b810c03d02fa404d8bdace84be8ccf99

Shannon, P. *et al*. 2003. Visualization Software Cytoscape. doi:

10.1101/gr.1239303. Retrieved January, 2021. {*Code and/or software*.}

Bolyen, E. *et al*. 2019 *Nat. Biotech*. QIIME 2. https://doi.org/10.1038/s41587-019-0190-3. Retrieved January 2021. {*Code and/or software*.}

## Results

### Urine is Rich in Metabolites

Untargeted metabolomics data was acquired and analyzed for 240 urine samples (4 storage conditions, +/-AssayAssure®,10 participants, in triplicate). Overall, the entire dataset generated 6,259 MS2 features using the GNPS MZmine2 workflow and 274 library matches through GNPS FBMN. 3,303 metabolites were networked to at least one other metabolite. 747 metabolite networks were identified which ranged from 2 to 94 metabolites in size. Of the 747 molecular families identified, all but two contained metabolites found in both the presence and absence of AssayAssure®.

### Individual participants drive data clustering while storage conditions have minimal effect

Feature finding in the non-AssayAssure® (nonAA) dataset resulted in fewer MS2 features (4,205 metabolites) and annotations (187 library matches) than the combined data set. Of these 2075 metabolites were networked to at least one other metabolite, resulting in 434 molecular families ranging from 2 to 99 metabolites in size. The beta diversity of the nonAA samples revealed a tight clustering by Participant ID on the Bray Curtis PCoA (Figure 1A). PERMANOVA analysis on the separation between participants showed a significant difference in the composition of the metabolomics data between participants (p-value = 0.001, pseudo-F = 64.013).

**Figure 1.**
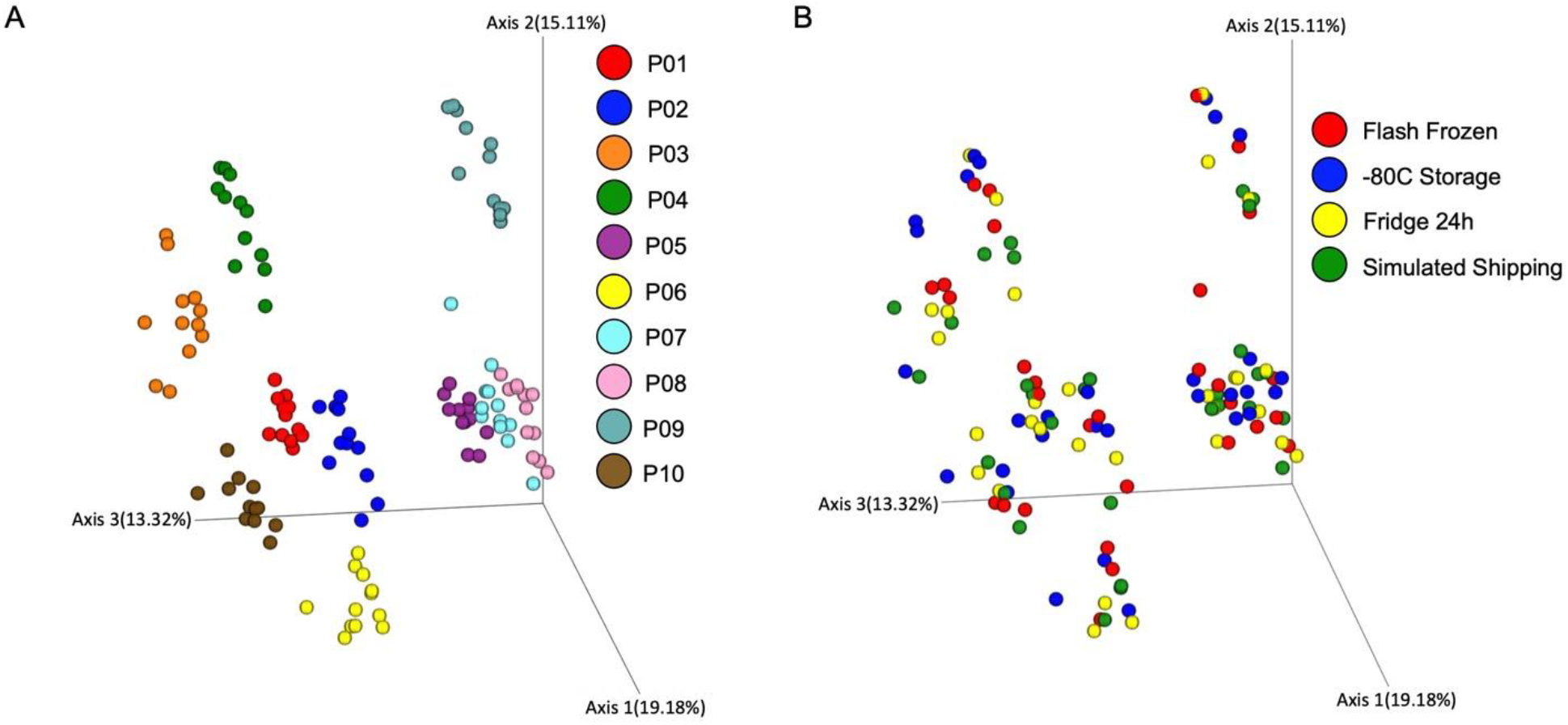
Urinary Metabolomic Profile by Participant and Storage Condition. A. Principal component analysis (PCoA) calculated using a Bray Curtis distance metric of metabolomics samples without AssayAssure®, colored by participant ID showing significant differences in metabolite composition between participants (PERMANOVA p = 0.001, pseudo-f = 64.013). B. Bray Curtis PCoA of metabolomics samples without AssayAssure®, now colored by storage method (flash frozen (red), freezer storage (blue), fridge 24h (yellow), simulated shipping (green)), showing no significant difference between storage conditions (PERMANOVA p = 0.999, pseudo-f = 0.359828).

When analyzed by storage condition, no separation or trends were seen due to storage condition on the PCoA plot (Fig 1B). PERMANOVA analysis also indicated that there was no significant clustering by storage condition (p-value = 0.999, pseudo-F = 0.359828). Random forest analysis using the QIIME 2 sample-classifier function performed slightly over baseline (accuracy ratio = 1.556), producing an ROC curve result of AUC=0.65. This performance was mainly due to the fact that the algorithm was able to predict the classification of the 3 month freezer samples with 100% accuracy. This high accuracy may indicate changes in abundance differences of metabolites when frozen for extended periods of time; however, changes were not large enough to see a significant difference between samples’ overall beta diversity, as seen with the non-significant PERMANOVA score.

As urine metabolite composition varied significantly amongst participants, we next examined separate participant tables in order to analyze the difference in storage conditions for each participant. Individual PCoAs of nonAA samples did not show distinct storage method separation (Example PCoA in Supplemental Figure 1A) and all participant level PERMANOVA analyses of nonAA samples resulted in insignificant variation between storage methods for every participant (Supplemental Figure 1B). Overall, for the samples not stored in AssayAssure®, there were no significant differences in metabolomic profiles between the four tested storage conditions (Flash frozen, 3 months in freezer, 24 hours in fridge or shipping simulation).

### Sample metabolomic trends are similar with and without AssayAssure®

Next, samples stored in AssayAssure® were analyzed. The GNPS MZmine workflow found 5,303 MS2 features and GNPS FBMN identified 238 library matches within these MS2 features. 2640 metabolites were networked to at least one other metabolite, creating 564 networks ranging in size from 2 to 88 metabolites. Overall, the AssayAssure® samples (AA) resulted in 1098 more MS2 features found by the MZmine software and 51 more annotations than the same samples stored without AssayAssure®. These results may indicate the presence of AssayAssure® contaminants in the data, as well as additional adducts and mass shifts.

However, the overall metabolomic profile of the data remained similar to what was seen with the nonAA samples. Beta diversity analysis of the data indicated a tight clustering by participant and no obvious trends from storage conditions, as visualized by PCoA (Supplemental Figure 2A and 2B). PERMANOVA analysis by Participant ID resulted in a significant differentiation between participants (p = 0.001, pseudo-f = 22.4337). PERMANOVA test of the storage conditions indicated no significant difference between the stored conditions across all participants (p-value = 0.975, pseudo-f = 0.573503). Additionally, the random forest analysis showed overall predictive accuracy of 0.694444, which is 2.77778 times the baseline accuracy. The algorithm was able to classify the test samples from the flash frozen group with 100% accuracy, and samples from the freezer 3-month group with 88% accuracy. Overall, this indicates that for samples with AssayAssure® the type of storage may slightly alter the preservation of certain metabolites. However, like the nonAA samples, the machine learning predictive ability does not result from an overall significant beta diversity profile shift of the data, as was demonstrated by the PERMANOVA test.

Next, we examined separate participant data to ascertain whether AssayAssure® differentially impacted metabolite composition by storage condition. Unlike the nonAA samples, samples from half of participants did show significant separation between sample storage conditions as shown in the example PCoA plot and in the table (Supplemental Figure 1C and 1D). To further assess whether this significant difference was secondary to metabolite abundance or presence/absence we calculated a beta diversity matrix using Jaccard: only 2 participants showed significant separation in the beta diversity of their samples by storage conditions (P=0.028 and 0.038) while pseudo-F values remained comparable to participants with non-significant metabolome separation (pseudo-F=1.22806 and 1.15052). Overall, this indicates subtle alteration in metabolite abundance based on storage conditions in the presence of AssayAssure®. The biologic variation between participants superseded the impact of AssayAssure® and storage conditions similar to nonAA samples.

### AssayAssure®shifts entire metabolome profile in a predictable way

We next examined how AssayAssure® influences metabolite composition in all samples. The metabolomic composition between AA samples and nonAA samples was visualized by a PCoA plot using a Bray Curtis distance metric (Figure 2A). Samples showed significant separation between those with AssayAssure® versus those without (p-value = 0.001, pseudo-F = 18.8938 by PERMANOVA). Of the 6,259 MS2 spectra in the entire dataset, 6 were unique to nonAA samples and 65 were unique to AssayAssure®. However, of the 747 molecular families identified the entire dataset, only 2 doublet networks were unique to AssayAssure®. The remaining unique metabolites (to AssayAssure®) were networked to metabolites present in nonAA samples. These data indicate that, although unique metabolites exist in the presence of AssayAssure®, these spectra represent modifications to metabolites already present in nonAA samples.

To determine the impact of AssayAssure® on metabolite abundance, a random forest analysis was performed. The model was able to accurately predict between nonAA and AA samples (accuracy ratio = 1.91667). To visualize how AssayAssure® changed the metabolite abundance, features we considered important from this learning model were annotated in the molecular network (Fig. 2B). The variable presence of significant features across metabolite networks indicates that metabolite networks were differentially impacted by the presence of AssayAssure®.

To explore the phenomena, we examined one of the largest molecular families containing a majority of the carnitine library matches, a common and biologically relevant urine metabolite^13,14^. A Dunn’s test was performed across the molecular family and the p-values comparing AA to nonAA abundance were visualized (Figure 3A). Supplemental Table 1 contains a list of every multiple test corrected p-value along with which group (AA vs nonAA) contained the highest median abundance of the feature. Overall, in this family 55 of the 85 features were significantly different between the AA versus nonAA samples. Forty-nine of these 55 features were of higher abundance when stored in AssayAssure®. In addition to the carnitine library matches within this molecular family, there were also a few potential bile acid, which were also significantly more abundant in the AA samples. Although the median abundance levels for many features were higher in the AA samples, these features were still present in the nonAA samples. The addition of AssayAssure® to the samples slightly altered the preferential preservation of some metabolites but did not completely add or completely lose any features in this molecular family.

**Figure 2.**
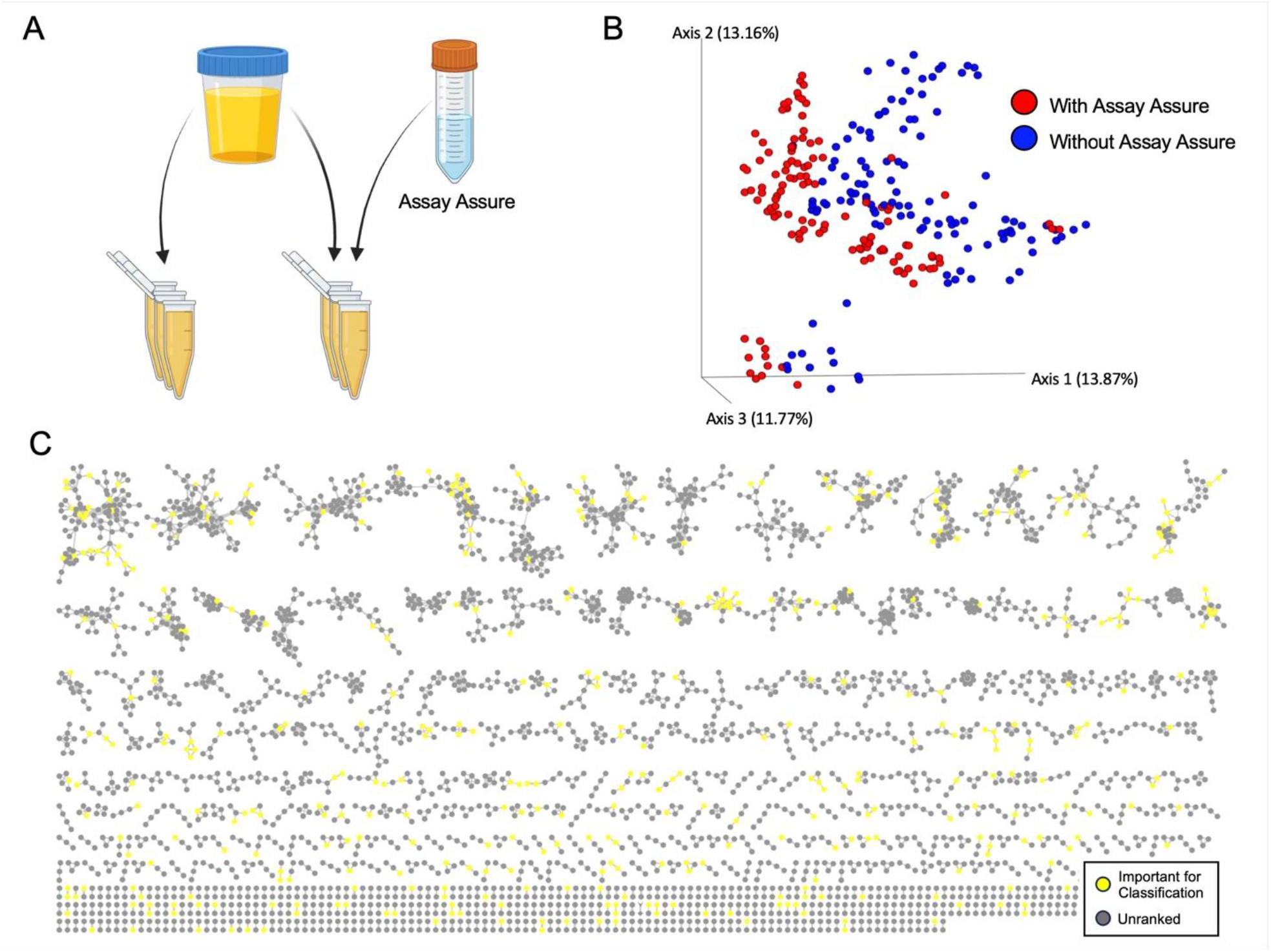
Urinary Metabolomic Profile in Presence/Absence of AssayAssure®. A. Samples were either stored with or without the addition of DNA preservative, AssayAssure®. B. Principal component analysis (PCoA) of metabolomics Bray Curtis of samples with (red) and without (blue) AssayAssure® (PERMANOVA p = 0.001, pseudo-f = 18.8938). B. Urinary metabolite GNPS network. Each circle represents a cluster of matches MS2 spectra (features), connected by edges where the width of the line indicates a higher cosine score for spectral alignment (minimum cosine = 0.70). Random forest importance score was used for coloration of clusters, with red indicating an importance score of 0.001 or higher.

**Figure 3.**
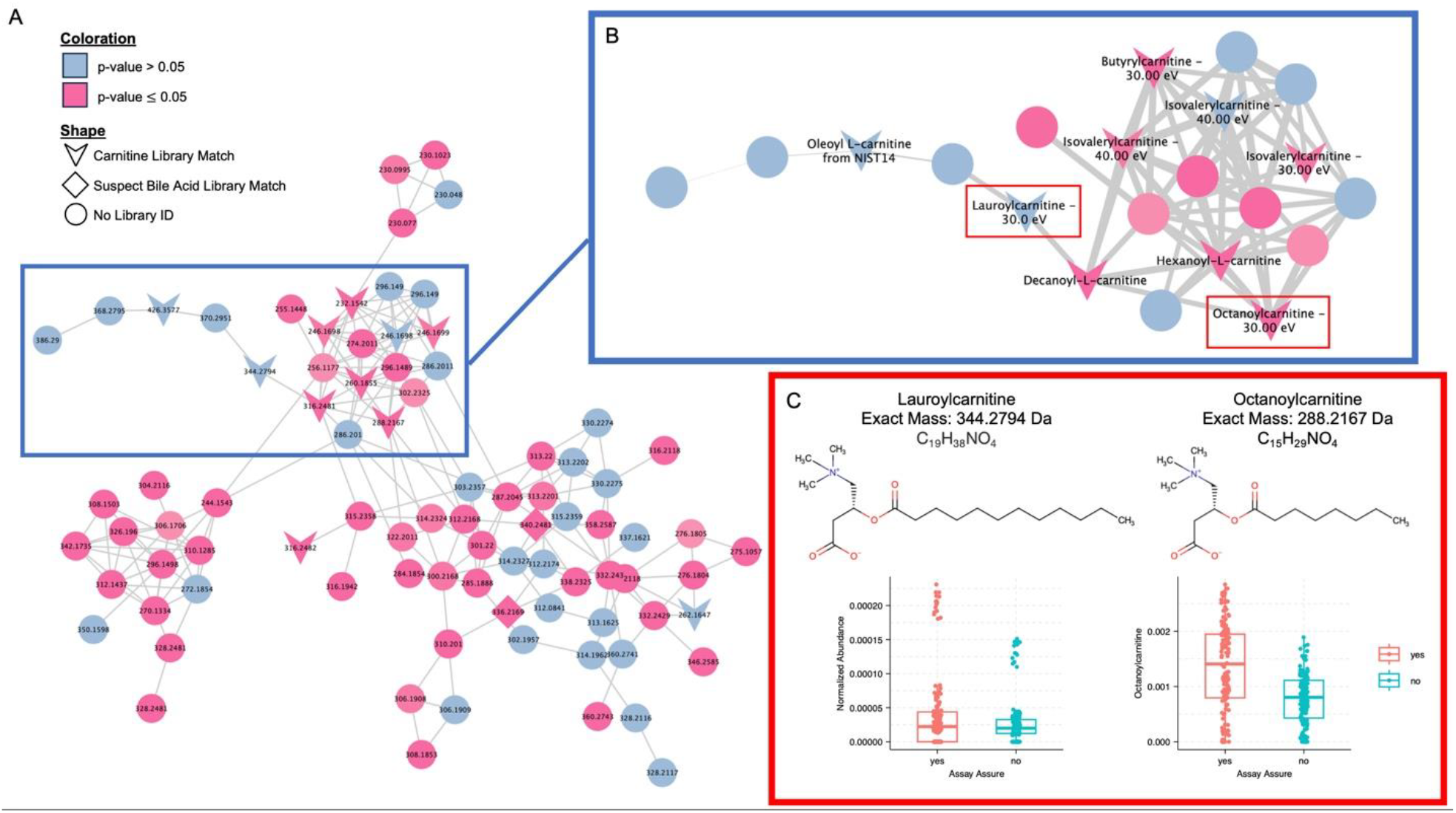
Carnitines are Differentially Preserved in AssayAssure®. A. The full carnitine molecular family network produced by GNPS FBMN. Each node represents a cluster of aligned spectra, identified as one feature. Nodes are labeled by parent mass (Da), shaped by library annotation status, and colored by a Dunn’s test conducted on the normalized abundance levels between AssayAssure® samples and those without it. Edges between nodes indicate a spectral connection, with a cosine score >0.70. B. A zoomed in portion of the larger molecular family (indicated by the blue boxes) where the nodes are now labeled by GNPS Library ID match. Shapes are still indicative of spectral match category; coloration is still by Dunn’s test. The widths of the edges are now indicative of a larger cosine score, with the larger edges being a cosine closer to 1. C. Two example metabolites from the network, indicated by the red boxes. Structures are presented and the normalized abundance levels are plotted. As with most of the carnitine metabolites, these example metabolites are found in higher abundance in the AssayAssure® samples

To determine whether the addition of AssayAssure® reduced interparticipant metabolome separation, the entire dataset was analyzed by participant. PERMANOVA test still found significant variation between individuals (p-value = 0.001, pseudo-F = 40.6167), despite the separation seen between AssayAssure® and nonAA. All participants can be distinguished from each other using their urine metabolome whether their samples were stored in the presence or absence of AssayAssure®. Additionally, storage condition was analyzed at this scale, and conditions were not significantly distinct from each other using the PERMANOVA analysis (p-value = 0.755, pseudo-F = 0.840106). Overall, the entire dataset is overwhelmingly separated by participant, and then within each participant cluster there is a separation by whether the samples were stored in the presence of AssayAssure®. Other storage conditions had no significant effect on separation when analyzed within the entire dataset.

## Discussion

Broad untargeted metabolomics projects are subject to batch effects due to the collection methods and sensitivity of the data type^15^. Consequently, it is unclear whether metabolomic analysis of stored and preserved urine would yield biologically relevant data. Here, we assessed whether common storage conditions in the presence or absence of the DNA preservative AssayAssure® impacted urine metabolomic profiles. There are two principal findings of this work: 1) the storage conditions tested do not significantly impact urine metabolite composition and 2) the presence of AssayAssure® is not associated with metabolite loss despite significant changes in abundance of some urinary metabolites. In addition, unique metabolites added by AssayAssure® network with metabolites that are seen in urine samples without AssayAssure®.

This analysis indicated that the effects of the four common storage conditions (refrigeration (2-4 h), simulated shipping on wet ice, frozen at -80°C (3 months) and quick-frozen urine) were minimal to the overall metabolomic profile. Metabolomic profile diversity in both samples with and without AssayAssure® did not vary by these four storage conditions. The machine learning analysis was able to learn and predict samples into their storage conditions above a baseline random approach, although this separation was not reflected in the overall beta diversity. This finding is consistent with the conclusion that storage conditions did not change the overall urinary metabolite profile of the samples, although the storage conditions may have resulted in the minor differential preservation of a few compounds in which the machine learning was able to identify and predict from. Analysis separated by participant showed no differences between storage conditions, indicating biologically relevant interparticipant variability outweighs the impact of storage conditions or the presence or absence of AssayAssure®.

The separation between the metabolomics data for sample stored with vs. without AssayAssure® is significant and not surprising, given the certainty that the preservative solution adds unique features. Interestingly though, this separation is not just due to the presence of AssayAssure® contamination features, but an overall shift in a large amount of feature abundances across the dataset, as seen with the machine learning visualization of the network. AssayAssure® is used to protect RNA and DNA from degradation within samples by suppressing enzymatic activity, which resulted in increased abundance of some metabolites as well. We illustrated metabolite compositional changes within the carnitine molecular family (Figure 3 and Supplemental Table 1). Notably, although important metabolites were not lost when using AssayAssure®, they were present in different abundances and adducts within networks.

Additionally, within the AssayAssure® samples there was more separation between the 4 storage conditions than when analyzing the non-AssayAssure® samples. Five of the 10 participant individual datasets indicated a significant separation between the 4 sample storage conditions. Caution is needed in interpretation, as this small sample sizes has the potential for skewed results. These data suggest preservation in the presence AssayAssure® may be dependent on storage conditions. Despite this variability, the interparticipant variability continued to supersede the impact of storage conditions or AssayAssure® presence. Overall, these data suggest that ideal sample storage and preservative conditions should be matched between participants. It also suggests that the tested conditions do not preclude metabolomic analysis. Finally, caution is recommended when interpreting subtle changes in metabolic abundance longitudinally or with variable storage for a given participant.

In all storage conditions analyzed in the presence or absence of AssayAssure®, biological metabolites were identified, and biological differences were still significant – indicating that samples from all conditions can be used for metabolomic analysis producing meaningful results. AssayAssure® is not a background signal; it results in differential preservation of metabolites as well as different adduct formation. Parent molecules are still present in all samples, indicating that it is plausible to use urine samples preserved with AssayAssure® for metabolomics analysis. This finding suggests biobank samples may be informative for both sequencing and metabolomics (multi-omics) projects.

Strengths of this work are that frequently employed storage and preservative conditions were studied making this analysis applicable to bio banked urine specimens stored under similar conditions. Additionally, this analysis included global profiling of metabolites rather than a targeted approach and is applicable for a wide range of clinically applicable metabolites.

Limitations of this data include the small samples size with a limited participant population that lacks phenotypic or demographic characterization of participants. Furthermore, only a single collection method: midstream, voided urine, was used, which is presumed to represent the urogenital and not urinary only metabolome. Additional studies are needed to determine the relationship between the voided urine metabolome and samples collected using other methods (ie. catheterized, suprapubic aspirate, etc.). Metabolites in lower abundance in participant populations or exclusively found in specific participant populations may not be represented in the metabolomic profiles used in this work. Consequently, these metabolites may have a divergent response to the conditions and preservative used in this study. In addition, we did not include prolonged frozen timepoint which may impact metabolite preservation.

## Supplemental Figures/Tables

**Supplemental Figure 1.**
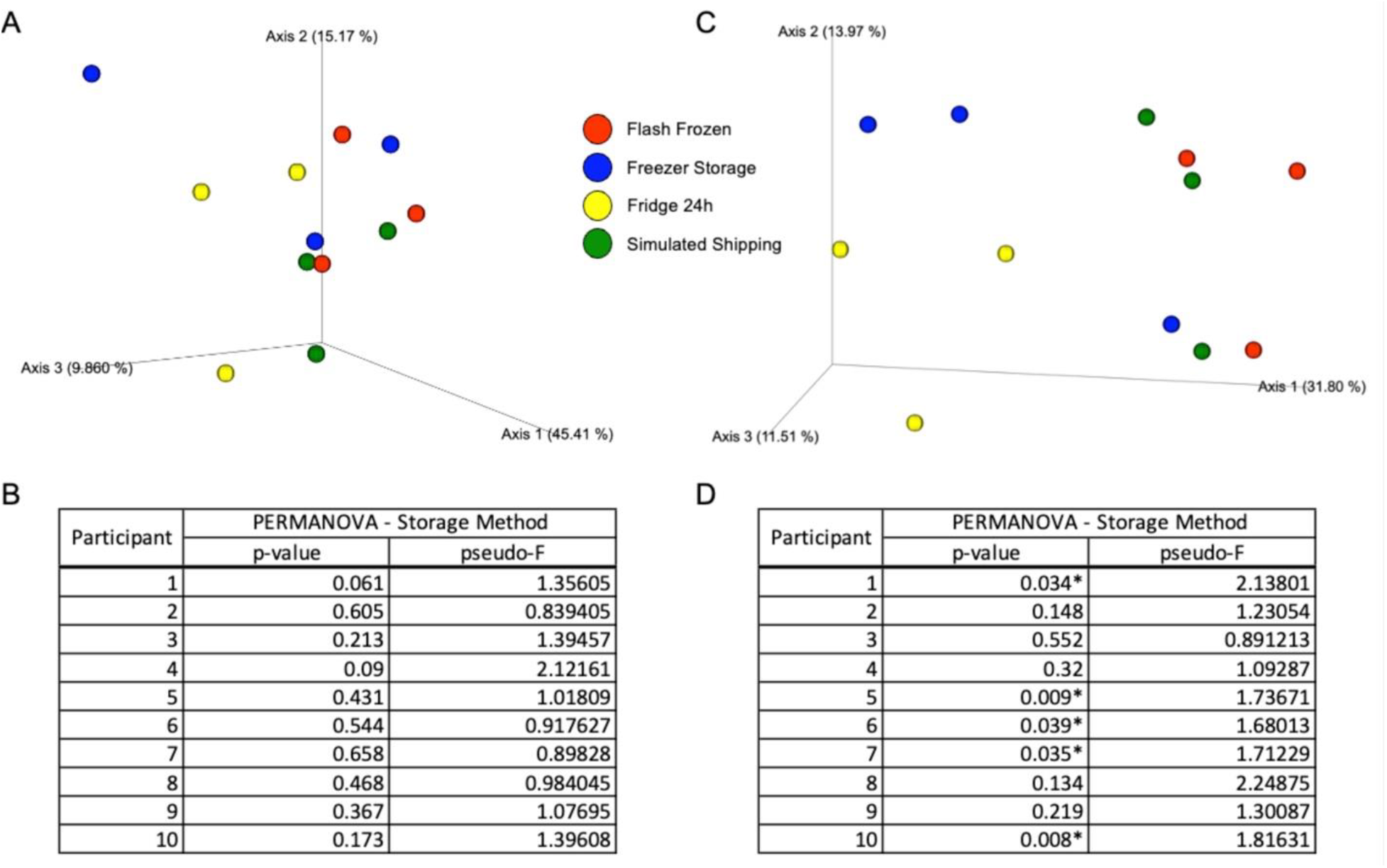
Example of individual participant storage condition analysis, using Participant 10 data. A. Bray Curtis PCoA plot of Participant 10’s samples stored without AssayAssure®, colored by storage condition. B. Table of the PERMANOVA scores for the samples without AssayAssure® for each individual participant, calculated from the Bray Curtis distance metric. No significant separation between storage condition was seen for any participant. C. Bray Curtis PCoA plot of Participant 10’s samples stored in AssayAssure®, colored by storage condition. B. Table of the PERMANOVA scores for the samples in AssayAssure® for each individual participant, calculated from the Bray Curtis distance metric. Significant variance between storage conditions can be seen 5/10 of the participants for the AssayAssure® stored samples.

**Supplemental Figure 2.**
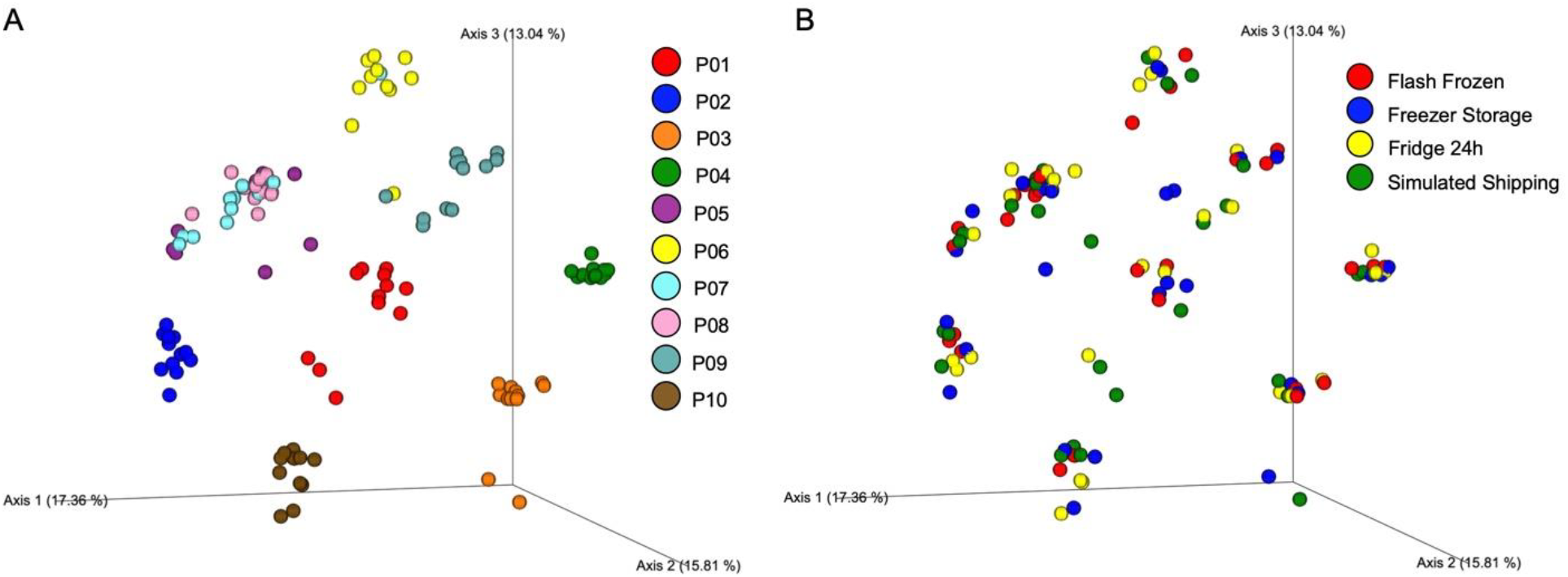
AssayAssure® stored Urinary Metabolomic Profile mimics samples without AssayAssure®. A. Principal component analysis (PCoA) calculated using a Bray Curtis distance metric of metabolomics samples stored in AssayAssure®, colored by participant ID showing significant differences in metabolite composition between participants (p = 0.001, pseudo-f = 22.4337). B. Bray Curtis PCoA of metabolomics samples stored in AssayAssure®, now colored by storage method (flash frozen (red), freezer storage (blue), fridge 24h (yellow), simulated shipping (green)), showing no significant difference between storage conditions (p = 0.999, pseudo-f = 0.359828).

**Supplemental Table 1.** Significant differences in abundance levels seen within the carnitine molecular family. This table contains every feature in the network visualized in Figure 3, labeled by feature ID or library ID if present. The p-values were calculated using a Dunn’s test and highlighted in red if significant (p < 0.05). The final column indicated in which sample type the median of the data is higher, a representation of which sample type the feature is more abundant in.

| Feature ID | AssayAssure p-value | Increased Median |
| --- | --- | --- |
| 326 | 0.832706 | Assay Assure |
| Butyrylcarnitine - 30.00 eV | 0.002702 | Assay Assure |
| 340 | 0.000013 | Basic Storage |
| 355 | 0.013439 | Assay Assure |
| 357 | 0.033795 | Assay Assure |
| (2R)-3-Hydroxyisovalerylcarnitine | 0.059809 | Basic Storage |
| 410 | 1.57E-08 | Assay Assure |
| Octanoylcarnitine - 30.00 eV | 3.09E-10 | Assay Assure |
| 439 | 0.019691 | Assay Assure |
| Isovalerylcarnitine - 40.00 eV | 0.188137 | Assay Assure |
| 515 | 0.960983 | Basic Storage |
| 520 | 0.000489 | Assay Assure |
| 580 | 0.855734 | Basic Storage |
| 622 | 0.000418 | Basic Storage |
| 636 | 9.54E-08 | Assay Assure |
| 741 | 0.144189 | Basic Storage |
| 756 | 0.012991 | Assay Assure |
| 760 | 0.000045 | Assay Assure |
| 769 | 0.034116 | Basic Storage |
| 772 | 2.24E-08 | Assay Assure |
| Candidate Annotation to Trihydroxylated bile acid | 5.46E-19 | Assay Assure |
| 809 | 1.19E-07 | Assay Assure |
| 817 | 0.000009 | Assay Assure |
| 827 | 6.31E-10 | Assay Assure |
| 857 | 0.000019 | Assay Assure |
| Decanoyl-L-carnitine | 0.000220 | Assay Assure |
| 886 | 0.114584 | N/A |
| 914 | 1.50E-07 | Assay Assure |
| 998 | 0.000694 | Assay Assure |
| Isovalerylcarnitine - 30.00 eV | 0.014045 | N/A |
| Hexanoyl-L-carnitine | 0.000120 | Assay Assure |
| 1091 | 0.001096 | Assay Assure |
| 1223 | 0.303105 | Basic Storage |
| 1239 | 0.000326 | Assay Assure |
| 1272 | 0.000025 | Assay Assure |
| 1323 | 5.62E-10 | Assay Assure |
| 1335 | 0.000144 | Assay Assure |
| 1378 | 0.000895 | Assay Assure |
| Candidate Annotation to Dihydroxylated bile acid | 2.96E-13 | Assay Assure |
| 1397 | 0.000467 | Assay Assure |
| 1507 | 0.001902 | Assay Assure |
| 1602 | 0.003471 | Assay Assure |
| Decanoyl-L-carnitine | 0.000170 | Assay Assure |
| 1710 | 0.397258 | N/A |
| Lauroylcarnitine -30.00 eV | 0.294719 | Assay Assure |
| 2029 | 0.000097 | Assay Assure |
| 2044 | 0.063070 | Basic Storage |
| 2052 | 0.171425 | Assay Assure |
| 2090 | 0.000073 | Assay Assure |
| 2356 | 0.035581 | Assay Assure |
| 2501 | 0.336170 | Basic Storage |
| 2521 | 0.962024 | N/A |
| 2706 | 0.000012 | Assay Assure |
| 2931 | 0.091564 | Assay Assure |
| 3174 | 0.385692 | N/A |
| 3232 | 0.000071 | Assay Assure |
| 3249 | 3.37E-09 | Assay Assure |
| 3745 | 0.000004 | Assay Assure |
| 3764 | 0.582465 | Basic Storage |
| 3773 | 1.78E-11 | Assay Assure |
| 3874 | 0.000032 | Assay Assure |
| 4059 | 0.000000 | Assay Assure |
| Isovalerylcarnitine - 40.00 eV | 0.015503 | Assay Assure |
| 4217 | 0.068954 | N/A |
| 4749 | 0.017391 | Assay Assure |
| 5094 | 0.639176 | N/A |
| 5396 | 0.015746 | Assay Assure |
| 5463 | 0.882714 | Assay Assure |
| 5558 | 0.017178 | Assay Assure |
| 5581 | 0.000645 | Assay Assure |
| 5583 | 0.000107 | Assay Assure |
| 5821 | 0.243195 | Basic Storage |
| 6353 | 0.460730 | N/A |
| 7343 | 0.723387 | N/A |
| 7562 | 0.392616 | N/A |
| 8033 | 0.000028 | Assay Assure |
| 8465 | 0.959533 | Basic Storage |
| 8489 | 0.893419 | Basic Storage |
| 8564 | 0.037341 | N/A |
| 9132 | 0.061693 | Basic Storage |
| 9402 | 0.427908 | N/A |
| 9954 | 0.000135 | N/A |
| Spectral Match to Oleoyl L-carnitine from NIST | 0.143807 | N/A |
| 10568 | 0.001695 | Assay Assure |
| 11114 | 0.261542 | Assay Assure |

